# RNA degradation by ribonuclease T2 is required for phosphate homeostasis in *S. cerevisiae*

**DOI:** 10.64898/2026.07.20.739639

**Authors:** Allison M. Kee, Nusrat Jahan, Davion J. McFall, Cote Briggs, Lucas R. Girard, Samuel Spiese, Jennifer F. Garcia

**Affiliations:** School of Biological Sciences, College of Arts and Sciences University of New England 11 Hills Beach Road, Biddeford, Maine 04005; Department of Molecular Biology Colorado College 14 East Cache La Poudre, Colorado Springs, CO 80906; School of Molecular and Physical Sciences, College of Arts and Sciences University of New England 11 Hills Beach Road, Biddeford, Maine 04005

## Abstract

Inorganic phosphate (P_i_) is a critical building block for key biomolecules including ATP, DNA, RNA, and phospholipids. Consequently, cells monitor phosphate levels and acquire phosphate when intracellular levels become low. Here we demonstrate an unanticipated connection between the enzymatic activity of the *S. cerevisiae* RNase T2 ortholog, Rny1, and inorganic phosphate availability. Rny1 has been studied for its role in autophagy-linked RNA degradation under starvation conditions. Here we find that in nutrient-rich conditions, cells lacking Rny1 function exhibit phosphate starvation phenotypes and aberrantly activate the *PHO* signaling pathway despite the presence of high levels of inorganic phosphate in the growth media. This activation is evidenced by increased *PHO* gene transcript levels and increased nuclear localization of Pho4 in *rny1Δ* strains. Complementation of *rny1Δ* with wild-type *RNY1* and human RNase T2 restores *PHO* transcript levels to those typically observed in exponentially growing cells. This implicates RNase T2-dependent RNA degradation as required for maintaining intracellular phosphate levels, even under phosphate-replete conditions. Furthermore, consistent with its potential role in freeing phosphate from degraded RNA, *RNY1* expression is itself further induced under phosphate-limited conditions. These observations suggest that RNase T2-mediated RNA decay is a part of a potentially conserved metabolic recycling pathway that provides P_i_ to growing cells. These findings reframe RNA as a key metabolic resource and positions RNase T2 enzymes, and their function in RNA degradation, as an unexpected player in phosphate homeostasis.

**Graphical Abstract:** 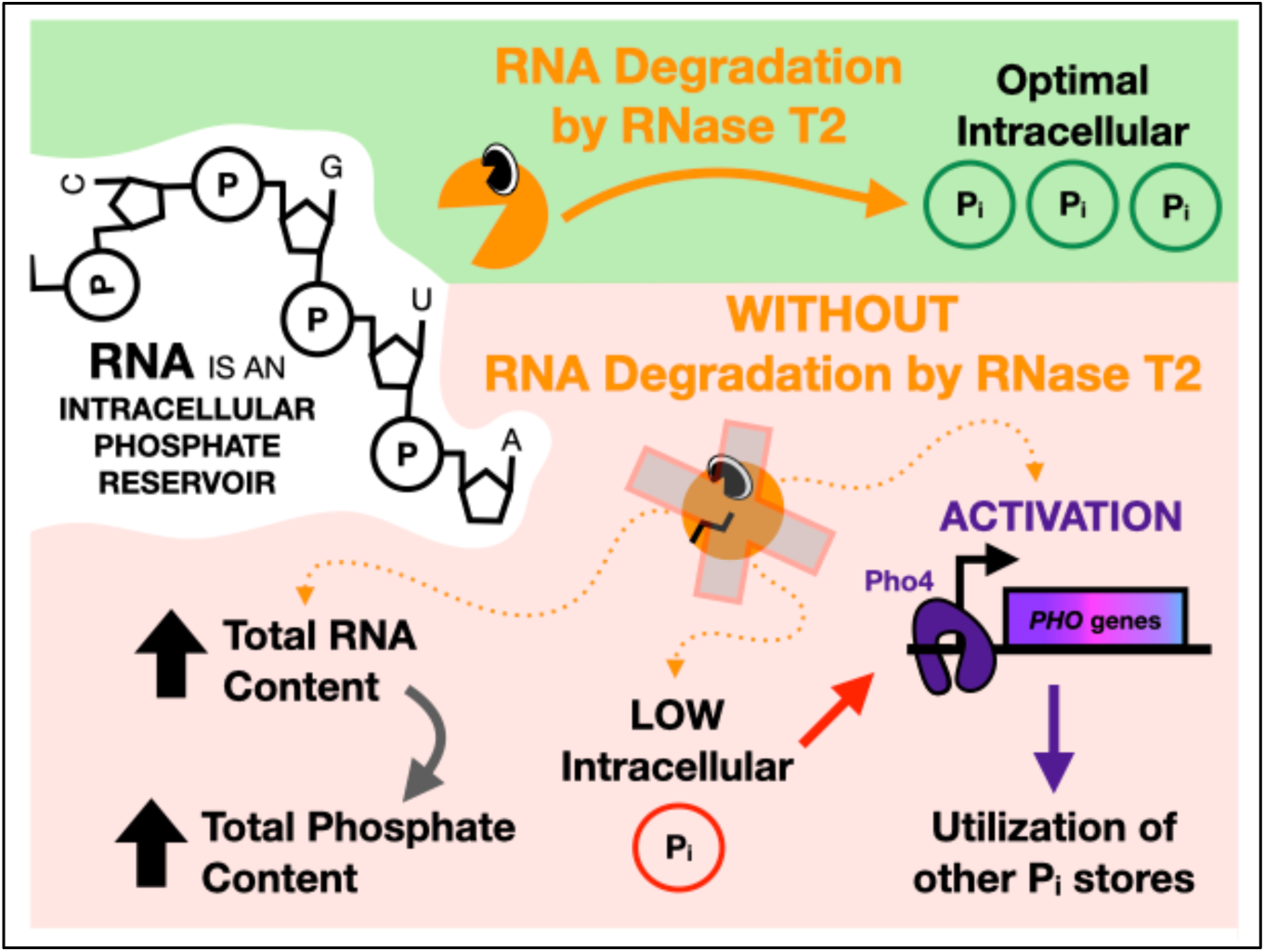

## Introduction

Conserved from bacteria to humans, RNase T2 enzymes form a group of ribonucleases present in a wide range of organisms. These enzymes possess endonucleolytic activity and cleave RNA into 3’-monophosphorylated nucleotides (MacIntosh 2011). RNase T2 family enzymes are typically localized to acidified organelles such as the lysosome in metazoans or the vacuole in yeast, where they carry out their function. RNase T2 enzymes have also been proposed to be secreted into the extracellular space (Acquati *et al*. 2011) or cytoplasm (Thompson and Parker 2009) to cleave RNA. RNase T2 enzymes have previously been characterized for their roles in stress induced RNA degradation by autophagy (Huang *et al*. 2015), particularly under conditions of nitrogen depletion (Huang *et al*. 2015, Makino *et al*. 2021) or oxidative damage (Thompson *et al*. 2008, Thompson and Parker 2009), suggesting they fulfill functions related to surviving metabolic stressors. Furthermore, RNase T2 enzymes have been implicated in tumor suppression (Acquati *et al*. 2011), neurodevelopmental disorders (Henneke *et al*. 2009, Haud *et al*. 2011), and immunity (Greulich *et al*. 2019), reflecting their broad relevance.

In *Saccharomyces cerevisiae*, the *RNY1* gene represents the sole RNase T2 homolog. Rny1 has been shown to contribute to the degradation of RNAs during stress conditions involving autophagy. Several studies have expanded our understanding of the role of Rny1 in cellular homeostasis. Minami *et al*. (2025) showed that Rny1 mediates selective autophagy of ribosomes during nitrogen starvation, facilitating rRNA degradation and contributing to the cellular response to nutrient stress. Indeed, autophagy in yeast selectively targets translating mRNAs for vacuolar degradation under rapamycin-induced stress (Makino *et al*. 2021). Another major class of RNAs targeted by Rny1 includes tRNAs, where degradation by Rny1 occurs under conditions of oxidative stress (Thompson and Parker 2009). This has a significant impact on translation, causing codon-specific pausing during oxidative stress (Pelechano, Wei, and Steinmetz 2015). These findings highlight the importance of Rny1 in RNA turnover during stresses such as nitrogen starvation and oxidative stress.

Here, we build upon these findings by expanding the role of RNase T2 enzymes to inorganic phosphate acquisition and the cellular stress of phosphate limitation. RNA can serve as a substantial intracellular phosphate reservoir, comprising an estimated 7-12% of the dry weight of a yeast cell (Delgado *et al*. 2013). Most of this pool (∼80-85% of the total RNA) consists of ribosomal RNA (rRNA), which is a known target of RNase T2 degradation via autophagic RNA decay (Haud *et al*. 2011, Hillwig *et al*. 2011, Huang *et al*. 2015, Makino *et al*. 2021). Under stress conditions such as nitrogen starvation, RNAs are non-selectively targeted to the vacuole and cleaved into 3’-nucleotide monophosphates by Rny1. The vacuolar phosphatase, Pho8, subsequently hydrolyzes the phosphate group from these Rny1 decay products, allowing the resulting nucleosides to be excreted into the extracellular space (Huang *et al*. 2015). The cellular fate of the freed inorganic phosphate (P_i_) remains uncharacterized. Given the scale of the RNA pool and the importance of P_i_, we hypothesized that RNase T2-mediated RNA breakdown functions as a conserved metabolic recycling pathway to maintain intracellular phosphate homeostasis.

Inorganic phosphate is essential for numerous cellular functions, including nucleic acid synthesis, energy currency production, metabolism, and phospholipid formation. Because of this, high levels of intracellular P_i_ are required for proper cell function, growth, and division. Yeast cells utilize the *PHO* signaling pathway to maintain and monitor intracellular P_i_ levels. Under low-P_i_ conditions, the Pho85-Pho80 CDK complex is inhibited, enabling the phosphate-responsive transcription factor, Pho4, to translocate to the nucleus (Schneider, Smith, and O’Shea 1994, O’Neill *et al*. 1996). Pho4 binds to sites with a consensus sequence (CACGTG; Harbison *et al*. 2004) and, together with a second transcription factor, Pho2, forms a Pho4– Pho2 complex that activates genes involved in phosphate acquisition and utilization, collectively known as the *PHO* genes (Zhou and O’Shea 2011). Under conditions where intracellular P_i_ levels are high, Pho4 nuclear translocation is suppressed by Pho85-Pho80 CDK phosphorylation, thus preventing *PHO* pathway activation (O’Neill *et al*. 1996).

In *S. cerevisiae*, excess intracellular phosphate is stored predominantly in the vacuole as inorganic polyphosphate (PolyP_i_; Yang *et al*. 2017). PolyP_i_ is a linear polymer made of orthophosphate monomeric units linked by phosphoanhydride bonds, similar to those present in ATP. PolyP_i_ synthesis is catalyzed by the vacuolar transporter chaperone (VTC) complex, with Vtc4 being the key polyphosphate polymerase (Ogawa, DeRisi, and Brown 2000). Under phosphate limitation, PolyP_i_ serves as a phosphate reservoir that helps maintain intracellular phosphate levels, as the *PHO* pathway induces several enzymes that free Pi from polyphosphates in low-P_i_ conditions (Shi and Kornberg 2005).

Here we show that cells harboring a deletion of the *RNY1* gene (*rny1*Δ) aberrantly induce the *PHO* pathway in a Pho4-dependent manner during growth in media containing high levels of extracellular phosphate. This induction is associated with higher total cellular phosphate levels but reduced intracellular free phosphate and polyphosphate levels compared with wild-type cells. Furthermore, *rny1Δ* cells accumulate higher levels of total RNA, supporting a model in which RNA acts as a major intracellular phosphate reservoir that can be released by RNase T2 activity. In line with this model, *rny1Δ* phenotypes are rescued by expressing catalytically active yeast and human forms of RNase T2 but not the catalytically inactive forms that cannot degrade RNA. Finally, we observe that *RNY1* is expressed under conditions of phosphate limitation. Collectively, our data indicate that RNase T2-dependent RNA degradation serves to maintain cellular phosphate homeostasis by reclaiming these internal P_i_ molecules.

## Materials and Methods

### Yeast Growth and Sample Collection

From frozen yeast stocks, yeast strains were streaked out on selective SC media and grown on the plate at 25-30°C for 1-3 days until visible patches were apparent. Yeast were cultured in 3-10 mL liquid synthetic complete (SC) media and incubated overnight at 30°C with shaking. Following 16-20 hrs of growth, the optical density at 600 nm (OD_600_) using a spectrophotometer was determined and cultures were diluted with fresh SC medium to an OD_600_ of 0.2 and incubated at 30 °C with shaking until collection.

To collect samples for subsequent biochemical analysis, OD_600_ was measured again and each culture was transferred to sterile conical tubes. Cells were harvested by centrifugation at 5k × g for 2 min, the supernatant was removed, and cell pellets were immediately flash frozen in liquid nitrogen. Samples were stored at –80 °C until further processing.

### Strain construction

All yeast strains and plasmids used in this study are listed in Table 1 and Table 2, respectively. The primers used for strain construction are detailed in Table 3.

**Table 1:** Yeast strains.

| Strain ID | Genotype | Source |
| --- | --- | --- |
| SC 1 | *his3Δ1, leu2Δ0, met15Δ0, ura3Δ0* | Yeast Deletion Collection |
| JG 4 | *rny1Δ::KanMX, his3Δ1, leu2Δ0, met15Δ0, ura3Δ0* | Yeast Deletion Collection |
| SC 29 | *rny1Δ::KanMX, his3Δ1, leu2Δ0, met15Δ0, ura3Δ0* | This study |
| SC112 | *his3Δ1, leu2Δ0, met15Δ0, ura3Δ0 + pRS415* | This study |
| SC113 | *rny1Δ::KanMX, his3Δ1, leu2Δ0, met15Δ0, ura3Δ0 + pRS415* | This study |
| SC114 | *rny1Δ::KanMX, his3Δ1, leu2Δ0, met15Δ0, ura3Δ0 + pRS415-RNY1* | This study |
| SC115 | *rny1Δ::KanMX, his3Δ1, leu2Δ0, met15Δ0, ura3Δ0 + pRS415-rny1-ci* | This study |
| SC167 | *his3Δ1, leu2Δ0, met15Δ0, ura3Δ0 + pCM189* | This study |
| SC143 | *rny1Δ::KanMX, his3Δ1, leu2Δ0, met15Δ0, ura3Δ0 + pCM189* | This study |
| SC179 | *his3Δ1::pPHO84-mCherry-HIS3+, leu2Δ0, met15Δ0, ura3Δ0* | This study |
| SC180 | *his3Δ1::pPHO84-mCherry-HIS3+, rny1Δ::KanMX, leu2Δ0, met15Δ0, ura3Δ0* | This study |
| SC 177 | *pho4Δ::KanMX, his3Δ1, leu2Δ0, met15Δ0, ura3Δ0* | Yeast Deletion Collection |
| SC196 | *pho4Δ::KanMX, his3Δ1::pPHO84-mCherry-HIS3+, leu2Δ0, met15Δ0, ura3Δ0* | This study |
| SC199 | *rny1Δ::KanMX, pho4Δ::HphMX, his3Δ1::pPHO84-mCherry-HIS3+, leu2Δ0, met15Δ0, ura3Δ0* | This study |
| SC7 | *HTB2::HTB2-GFP-His3MX6, his3Δ1, leu2Δ0, met15Δ0, ura3Δ0* | Yeast GFP Collection |
| SC193.3 | *HTB2::HTB2-GFP-His3MX6, PHO4::PHO4-mCh-HygB, rny1Δ::KanMX, his3Δ1, leu2Δ0, met15Δ0, ura3Δ0* | This study |
| SC163 | *rny1Δ::kanMX, his3Δ1, leu2Δ0, met15Δ0, ura3Δ0 + pCM189-RNaseT2* | This study |

**Table 2:** Plasmids.

| Plasmid ID | Plasmid | Selection Marker | Source |
| --- | --- | --- | --- |
| B4 | pFA6a-KanMX | *AmpR* | Bähler et al. 1998 |
| B23 | pAG32 (HpHMX) | *AmpR* | Goldstein and McCusker 1999 |
| B60 | pRS415 | *AmpR*; yeast = *LEU2+* | Sikorski and Hieter 1989 |
| B100 | pCM189 | *AmpR*; yeast = *URA3+* | From Euroscarf, Plasmid # P30326 |
| B69 | pRS415-*RNY1* | *AmpR*; yeast = *LEU2+* | This study |
| B85 | pRS415-*rny1-ci* | *AmpR*; yeast = *LEU2+* | This study |
| B131 | pCM189-*RNaseT2* | *AmpR*; yeast = *URA3+* | This study |
| B145 | pSCPPX2 | *AmpR* | From Addgene, Plasmid #112877 |
| B65 | pRP1618 | *AmpR* | Thomspon and Parker 2009 |
| B61 | pRP1619 | *AmpR* | Thomspon and Parker 2009 |
| B88 | pRP1621 | *AmpR* | Thomspon and Parker 2009 |
| P1-B1 | pAG303-GPD-ccdb | *AmpR*; yeast = *HIS3+* | From Addgene, Plasmid # 14134 |
| B134 | pFA6a-mCherry-HphMX | *AmpR* | From Addgene, Plasmid # 105156 |
| B140 | p(Pho4-BamI-PmeI) | *AmpR* | This study |

**Table 3:**
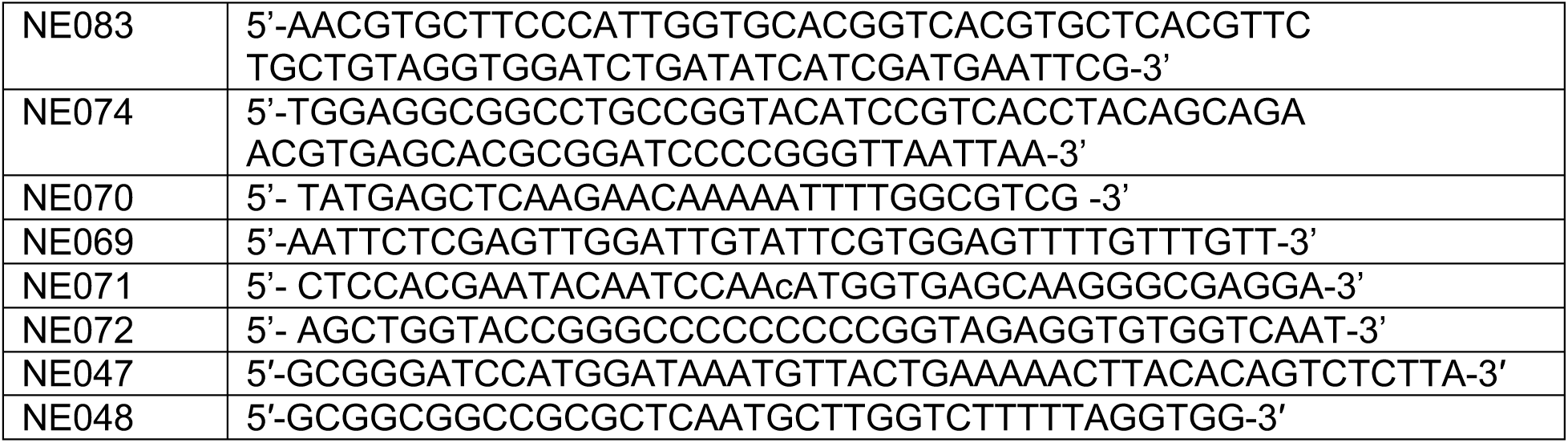
Primers used for strain construction.

Gene deletions (*rny1Δ* and *pho4Δ*) were generated via homologous recombination using PCR-amplified cassettes from pFA plasmids. Amplification utilized F1 and R1 primers containing 5′ overhangs homologous to the regions flanking the CDS. After selection, transformants were validated by colony PCR using gene specific primers.

To introduce a *Pho4-mCherry* fusion into yeast, the PHO4-mCherry-HphMX cassette was PCR-amplified from plasmid B134 (pFA6a-mcherry-hphMX) using primers NE083 and NE074. The resulting PCR product was transformed into the yeast GFP collection strain *HTB2-GFP-HIS3MX*. Transformants were selected on YPAD + Hygromycin B and integration was verified via colony PCR.

To construct pPHO84-mCh-tADH1, the *PHO84* promoter was PCR-amplified from BY4741 genomic DNA using primers NE070 and NE069. A 4.3-kb fragment from *Sac*I/*Xho*I-digested pAG303-GPD-ccdb was gel-purified and ligated with the *Sac*I/*Xho*I-digested *PHO84* PCR product to yield pAG303-pPHO84. An mCherry-tADH1 fragment was PCR-amplified from pFA6a-mCherry-HphMx using primers NE071 and NE072 and inserted into *Xho*I-digested pAG303-pPHO84 via Gibson assembly. The final B143 plasmid was linearized with *Nde*I and transformed into wild-type, *rny1Δ*, and *pho4Δ* strains to generate the p*PHO84-mCh* reporter strains. Integration of construct was validated growth on SC-HIS followed by colony PCR.

Plasmids pRS415-*RNY1* and pRS415-*rny1-ci* were constructed via restriction digest and ligation. Briefly, the pRS415 vector and donor plasmids (pRP1618 and pRP1619, respectively) were digested with *BamHI* and *SacI*. The resulting fragments were ligated, transformed into *E. coli*. Resulting plasmids were verified by Sanger sequencing.

To construct pCM189-*RNase T2*, the insert was PCR-amplified from pRP1621 using primers NE47 and NE48. The insert and pCM189 vector were digested with *BamHI* and *NotI*, joined via T4 DNA ligase, and verified by Sanger sequencing.

### RNA-Seq Library Preparation and High-Throughput Sequencing

Enriched poly(A)+ mRNA was isolated from 5 µg of intact total RNA using the NEBNext Poly(A) mRNA Magnetic Isolation Module (New England Biolabs). For library construction, the isolated mRNA fraction was immediately processed with the NEBNext Ultra II Directional RNA Library Prep Kit for Illumina (New England Biolabs). To generate a target insert profile of approximately 200 nt, purified mRNA was thermally fragmented at 94°C for exactly 8 min in the presence of divalent fragmentation parameters.

First-strand cDNA synthesis was initiated via random hexamer priming and executed in a thermocycler using sequential holds of 10 min at 25°C, 15 min at 42°C, and 15 min at 70°C. Second-strand cDNA synthesis was immediately performed at 16°C for 1 hr to incorporate dUTPs for strand directionality. Double-stranded cDNA fragments were purified using a 1.8X volume ratio of NEBNext Sample Purification Beads and eluted in 0.1X TE buffer.

Purified cDNA underwent end-repair and dA-tailing (30 min at 20°C followed by 30 min at 65°C) before undergoing adaptor ligation. Hairpin-loop NEBNext adaptors were prepared via a 25-fold dilution in Adaptor Dilution Buffer and ligated to the dA-tailed fragments for 15 min at 20°C, followed by a 15-minute incubation at 37°C with USER enzyme to excise the loop structures. The adaptor-ligated library fragments were purified using a 0.9X bead-to-sample ratio and enriched via targeted PCR multiplication using 13 amplification cycles.

Multiplexed library tracks were subsequently loaded onto an Illumina platform configured for Paired-End (PE) sequencing to yield structured 50 bp reads (2 × 50 bp).

### Differential Gene Expression Analysis

Differential gene expression analysis was performed using the UseGalaxy.org platform. Raw sequencing reads were first processed to remove adapter sequences and low-quality bases/reads using Trim Galore! (Galaxy Version 0.6.7). Quality-trimmed reads were mapped and counted using Sailfish (Galaxy Version 0.10.1.1) and the reference transcriptome (rna.fna) from the NCBI RefSeq assembly GCF_000146045.2 and reference genome of the *Saccharomyces cerevisiae* S288C genome build SGD R64-3-1.

Differential gene expression analysis was performed on the Sailfish count files using DESeq2 (Galaxy Version 2.11.40.7). Transcript-level abundances generated by Sailfish were aggregated to gene-level counts using tximport to evaluate differential gene expression comparing the *rny1*Δ count files against the WT count files as the baseline control. Size factors were calculated using the default median-of-ratios method, with gene-wise dispersions estimated using a parametric fit alongside standard DESeq2 independent filtering and Cook’s distance outlier detection under default parameters. Genes were classified as significantly differentially expressed if they exhibited an adjusted *p*-value < 0.05 and an absolute log₂ fold-change ≥ 1.

### Total RNA isolation

Total RNA was isolated from frozen yeast cell pellets using a modified acid phenol-chloroform extraction protocol (Roth, Madhani, and Garcia 2018). To minimize RNase contamination, all work surfaces and pipettes were decontaminated with RNase Away (Thermo Fisher Scientific). Liquid Nitrogen flash frozen cell pellets of 3-6 OD_600_ units were thawed on ice then washed with LET buffer (100mM LiCl; 10mM EDTA, pH 8.0; 10mM Tris-HCl, pH 7.5). The resulting cell pellet was resuspended in 150μL LET buffer and transferred to a 2ml shatter-resistant screw cap tube. Then 300μL of 0.5mm zirconia/silica beads and 150μL PL (phenol equilibrated with LET) was added. Samples were homogenized using a BeadBug 6 Microtube Homogenizer (Benchmark Scientific) for eight cycles of 30s at 2500 rpm. Following homogenization, samples were briefly centrifuged and 250μL RNase-free water was added. Lysates were centrifuged at 20k × g for 5min and the upper aqueous phase was transferred to a microcentrifuge tube containing Phase Lock Gel (PLG) and 250μL PCL (phenol–chloroform (1:1, v/v) equilibrated with LET). Samples were vortexed for 30s, and centrifuged at 20k × g for 5min. The aqueous phase was then transferred to a new PLG tube containing 400μL PCL, vortexed for 30s, and centrifuged again at 20k x g for 5min. The aqueous phase was then transferred to a new tube containing 400 μL chloroform and PLG, vortexed for 20s, and centrifuged at 20k x g for 5min. RNA was precipitated by transferring the final aqueous phase to a tube containing 40μL of 3M sodium acetate, followed by the addition of 1mL ice-cold 100% ethanol. Samples were mixed by inversion and incubated at –20°C for at least 1h. RNA was pelleted by centrifugation at 20k x g for 10 min at 4 °C. The supernatant was removed, and the pellet was washed with 500μL of ice cold 70% ethanol followed by centrifugation at 20k x g for 5 min at 4 °C. After removal of the ethanol wash, pellets were air-dried and resuspended in 100μL RNase Free water. RNA samples were stored at –20 °C.

### RNA Quality and Quantification for RT-qPCR

RNA concentration and purity were determined spectrophotometrically using a EzDrop spectrophotometer (Blue-Ray Biotech), Qubit RNA BR kit or Accublue RNA BR kit (Biotium). To assess RNA quality, 2µg of total RNA was run on a bleach gel as described in (Aranda et al. 2012).

### Preparation of Cell Lysates

Frozen cell pellets were thawed on ice and resuspended in cold Milli-Q water to obtain a final concentration of approximately 1 OD_600_ unit per mL. Two OD_600_ units was transferred to a 2mL screw-cap tube and pelleted by centrifugation at 5k x g for 2min. Cells were washed three times with 1 mL cold Milli-Q water, centrifuging at 5k x g for 2min between washes.

Following the final wash, pellets were resuspended in 200μL of breaking buffer (0.01% Triton X-100; 0.05% (w/v) SDS; 5mM NaCl; 0.5mM Tris-Cl, pH 8.0; 0.05mM EDTA, pH 8.0) and transferred to screw-cap tubes containing 0.5mm zirconia-silicate beads (Biospec). Cells were disrupted using a BeadBug 6 Microtube Homogenizer (Benchmark Scientific) for eight cycles of 30 s at 2.5k rpm. Lysates were transferred to fresh tubes and maintained on ice until further analysis.

### Total Protein Quantification of Cell Lysates

Total protein concentrations were determined using the Pierce Bicinchoninic Acid (BCA) Protein Assay Kit (Thermo Fisher Scientific) according to the manufacturer’s instructions with minor modifications. Cell lysates were diluted 1:5 in breaking buffer. A bovine serum albumin (BSA) standard curve was generated with concentrations ranging from 0 to 250μg/mL. Absorbance was measured at 562 nm using a SpectraMax MAX190 microplate reader. Protein concentrations were calculated from the BSA standard curve using a weighted line estimation in Microsoft Excel and corrected for dilution.

### Total RNA Quantification of Cell Lysates

Total RNA concentrations were measured using the AccuBlue Broad Range RNA Quantitation Kit (Biotium). Cell lysates were diluted in RNase-free water and assayed according to manufacturer’s instructions using the Qubit 4 fluorimeter. RNA standards ranging from 0 to 100ng/μL were prepared by serial dilution of the supplied RNA standard. RNA concentrations were calculated from raw RFU values using a weighted linear regression in Microsoft Excel, with adjustments made for sample dilutions.

### Determination of Free and Total Inorganic Phosphate in Cell Lysates

Free inorganic phosphate and total phosphate content were quantified using a malachite green colorimetric assay. For total phosphate measurements, cell lysates were mixed with an equal volume of 1N sulfuric acid and incubated at 100°C for 30 min to hydrolyze phosphate-containing metabolites, followed by cooling on ice. For free phosphate measurements, cell lysates were assayed without sulfuric acid treatment or heating. Phosphate concentrations were determined using a malachite green phosphate assay kit (MAK-307, Sigma-Aldrich) according to the manufacturer’s instructions. Absorbance was measured at 620nm using a SpectraMax MAX190 microplate reader, and concentrations were calculated using a standard curve generated from known phosphate standards.

### Purification of PPX

PPX was purified as described (Gray *et al*. 2014) with minor modifications. Chemically competent *E. coli* strains of BL21(DE3) cells were transformed with pSCPPX2. A single colony from frozen stock was cultured in Super Broth (Research Products International) with 100µg/mL Carbenicillin. After overnight growth at 37°C, the cells were induced with 1 mM IPTG and returned to the shaker at 37°C for 4hrs and then at 22°C for ∼20 hrs. Cells were then collected by centrifugation (4k x g for 20 min at 4°C) and stored at-80°C. Cells were re-suspended in 4 mL of Lysis Buffer (10 mM sodium phosphate; 0.1M NaCl; 1mM imidazole, pH 7.4; 50U/mL high salt tolerant Nuclease (NEB M0764); 1mg/mL Lysozyme (Sigma L6876); 1mM PMSF (Goldbio P-470) per gram of cell pellet and lysed using a Branson SFX150 tip sonicator (2 cycles of 2 min at 5 sec on, 5 sec off on ice). The cell lysate was collected from the supernatant following centrifugation at 15k x g for 20 min at 4°C. PPX purification was performed using column with Ni-NTA resin. The column was prepared with a binding buffer (50mM sodium phosphate, pH 8.0; 0.5M NaCl). After the cell lysate was added, the column was washed with Wash Buffer 1 (50mM sodium phosphate; 0.5M NaCl; 5mM imidazole; pH 8) and Wash Buffer 2 (50mM sodium phosphate, 0.5M NaCl, 20mM imidazole, pH 8) until flow-through did not detect protein using Bradford reagent. Elution of PPX was performed in elution buffer (50mM sodium phosphate; 0.5M NaCl; 0.5M imidazole, pH 8). Fractions containing PPX were pooled then concentrated using Pierce 10K MWCO Protein Concentrators. To remove phosphate and imidazole, concentrated PPX was dialyzed twice in dialysis buffer (20 mM Tris-HCl, pH 7.5; 50 mM KCl; 10% (v/v) glycerol). PPX was stored at 4 °C.

### Polyphosphate Isolation

Approximately 30 OD_600_ units of frozen cells were thawed on ice, washed with cold water then resuspended in 250µL LETS buffer (0.1M LiCl; 10mM EDTA; 10mM Tris, pH 8.0; 0.5% (w/v) SDS) and 250µL PL and homogenized with zirconia beads in a Bead-bug with eight 30 sec cycles at 2500 rpm then centrifuged (19k x g for 5 min at 4 °C) to separate the phases. The top aqueous layer was transferred into tubes with 500µL chloroform and phase-lock, briefly vortexed, and spun at 19k x g for 5 min. The top aqueous layer was then transferred to a new tube and precipitated with 2.5 volumes of cold 100% ethanol at-20°C overnight. Precipitated PolyP_i_ was then spun down at 19k x g for 5 min. After air drying, the pellets were resuspended in resuspension buffer (1mM EDTA; 10mM Tris-HCl, pH 8.0; 0.1% SDS) and their A260 measurements were taken on the EZdrop spectrophotometer to estimate PolyP_i_ content.

### Polyphosphate Quantification

To quantify polyphosphate (PolyP_i_), 8 A_260_ units of extracted sample were incubated in PPX reaction buffer (20 mM Tris-HCl, pH7.5; 5 mM MgCl2; 50 mM ammonium acetate) with 1.25 µg PPX1 at 37°C for 30 min. To determine total phosphate for normalization, an equivalent sample was hydrolyzed in 1N H_2_SO_4_ at 95°C for 30 min. Free phosphate in both reactions was quantified using the Malachite Green Phosphate Assay Kit as described above.

### Cell Fixation and Imaging

Cells were harvested by centrifugation and resuspended in 4% paraformaldehyde in 1x PBS. Following incubation at room temperature for 15 min, the cells were washed three times with 1x PBS and stored in 1x PBS at 4 °C until imaging. Imaging was performed using a Keyence BZ-X1000 microscope.

### Total RNA Staining

For total RNA staining, cells were harvested by centrifugation and washed twice with RNA staining preparation buffer (0.9M NaCl; 20mM Tris-HCl, pH 7.4). The cells were then incubated in RNA staining solution (0.9M NaCl; 20mM Tris-HCl, pH 7.4; 35% formamide; 500nM SYTO RNASelect Green, Invitrogen) for 30 min at 30 °C with shaking. Following incubation, the cells were washed twice with 1x PBS and fixed with ice-cold 100% methanol for 10 min on ice. Finally, the fixed cells were washed twice with 1× PBS and stored in 1× PBS at 4 °C until imaging.

### Microscopy data analysis

Pho84 reported gene expression was quantified from a max projection from a 4 slice substack centered on the most focus slices as determined by using the Find Focus Slices plugin. Cells were defined as ROIs using Cellpose BIOP wrapper (cyto2 model, diameter =20) followed by manual curation to remove moving cells and localized brightspots. Prior to measurements of CH3 (red) intensity for Pho84 quantification, the max projection was background corrected using a rolling ball of 10 pixels.

To measure SYTO RNAselect staining, maximum intensity z-projections were used to segmented images using the Cellpose cyto2 model, followed by manual curation moving cells and artifacts. To account for variations in focus across the z-plane, the normalized variance (variance/mean) for each region of interest (ROI) across all slices was used to identify a single optimal focal plane. To filter out blurred cells, individual ROIs were discarded if their normalized variance at this consensus slice fell below 99% of their peak individual variance. To minimize background noise and isolate the target signal, the optimal focal slice was subjected to a rolling-ball background subtraction. To correct for autofluorescence, the mean intensity of an unstained control strain was subtracted from the image prior to extracting the final quantitative measurements from the Cellpose generated ROIs.

To quantify Pho4 cellular localization, a 5-slice substack was generated from raw Z-stacks using ImageJ, centered on the slice with the highest variance on the CH2 (green) channel as identified by the “Find Focused Slices” ImageJ plugin. Total cell ROIs were generated from CH3 (red) maximum intensity projections using Cellpose (cyto3 model, diameter = 20, excluded on edges), followed by manual curation. Nuclear ROIs were segmented from CH2 maximum projections via autothresholding and particle analysis. Following rolling-ball background subtraction (radius = 10 pixels) on the CH3 max projections, the mean intensity was quantified for each nuclear and cytoplasmic ROI. Individual nuclear ROIs were matched to their corresponding cytoplasm ROI by utilizing the Cellpose mask files. To determine the localization ratio, the nuclear mean intensity was divided by the cytoplasmic mean intensity.

### RT-qPCR

Quantitative PCR (qPCR) was performed following the protocol of Roth *et al*. 2018. To remove contaminating genomic DNA, 10 µg of RNA was treated with 4 units of TurboDNase in 1x TurboDNase buffer for 45 min at room temperature. The DNase was inactivated by adding 0.5 M EDTA followed by heat treatment at 70 °C for 10 min. RNA was then purified using a Zymo Clean & Concentrator kit according to the manufacturer’s instructions. For cDNA synthesis and qPCR, Luna Supermix RT (New England Bio E3010) and Luna Universal qPCR Master Mix (New England Bio M3003) were used as substitutes for the reagents specified in the original protocol. To determine cDNA quantities, target C_q_ values were compared against a standard curve generated from genomic DNA of known concentrations. Sequences of primers used in RT-qPCR are listed in Table 4.

**Table 4:**
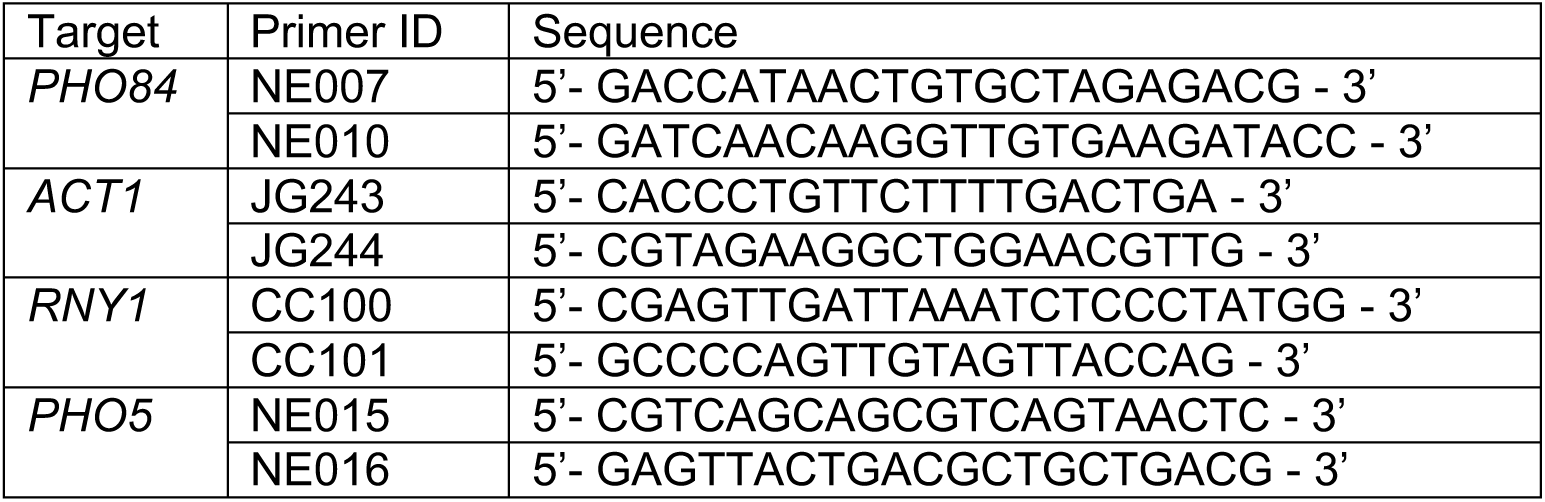
qPCR Primers.

## Results

### Cells lacking RNase T2 display aberrant Pho4-dependent PHO pathway activation

RNase T2 is a highly conserved enzyme that has been canonically studied under conditions of stress. However, given its evolutionary conservation from plants to humans and its critical roles in mammalian brain development, we sought to investigate its function during growth under standard nutrient-rich conditions. To determine if RNase T2 enzymes had key roles during periods of active growth and cell division, an undergraduate genomics lab course at Colorado College built RNA-seq libraries from poly(A) selected RNAs isolated from wild-type (WT) *S. cerevisiae* strains and yeast deletion collection strains that harbored a deletion of the sole yeast RNase T2 enzyme, *RNY1*. The resulting PE50 RNA-seq libraries were trimmed and aligned to the *S. cerevisiae* genome build R64-3-1. Differential gene expression analysis using DESeq2 revealed a subset of mRNAs that were upregulated by 2-fold or more in cells lacking RNase T2 function. This subset included 9 of the 80 Pho4-regulated mRNAs identified by Zhou and O’Shea (2011) as being induced during P_i_ starvation (Fig. 1A-B). Within this shared group, the core *PHO* genes, such as *PHO84*, *SPL2*, and *PHO5*, showed the most robust induction. Under low intracellular phosphate conditions, Pho4 undergoes nuclear translocation to activate the transcription of these *PHO* genes, which are required for external phosphate acquisition (e.g. *PHO84* and *PHO5*) or internal utilization of P_i_ stores (e.g. *VTC1* and *PPN2*). These results suggested that cells lacking RNase T2 function seemingly activate the *PHO* genes during growth in phosphate-rich media. This finding was unexpected, as the Synthetic Complete (SC) media used for these experiments contains high concentrations of P_i_ (∼0.7g/L), a condition not typically associated with *PHO* gene activation.

**Figure 1:**
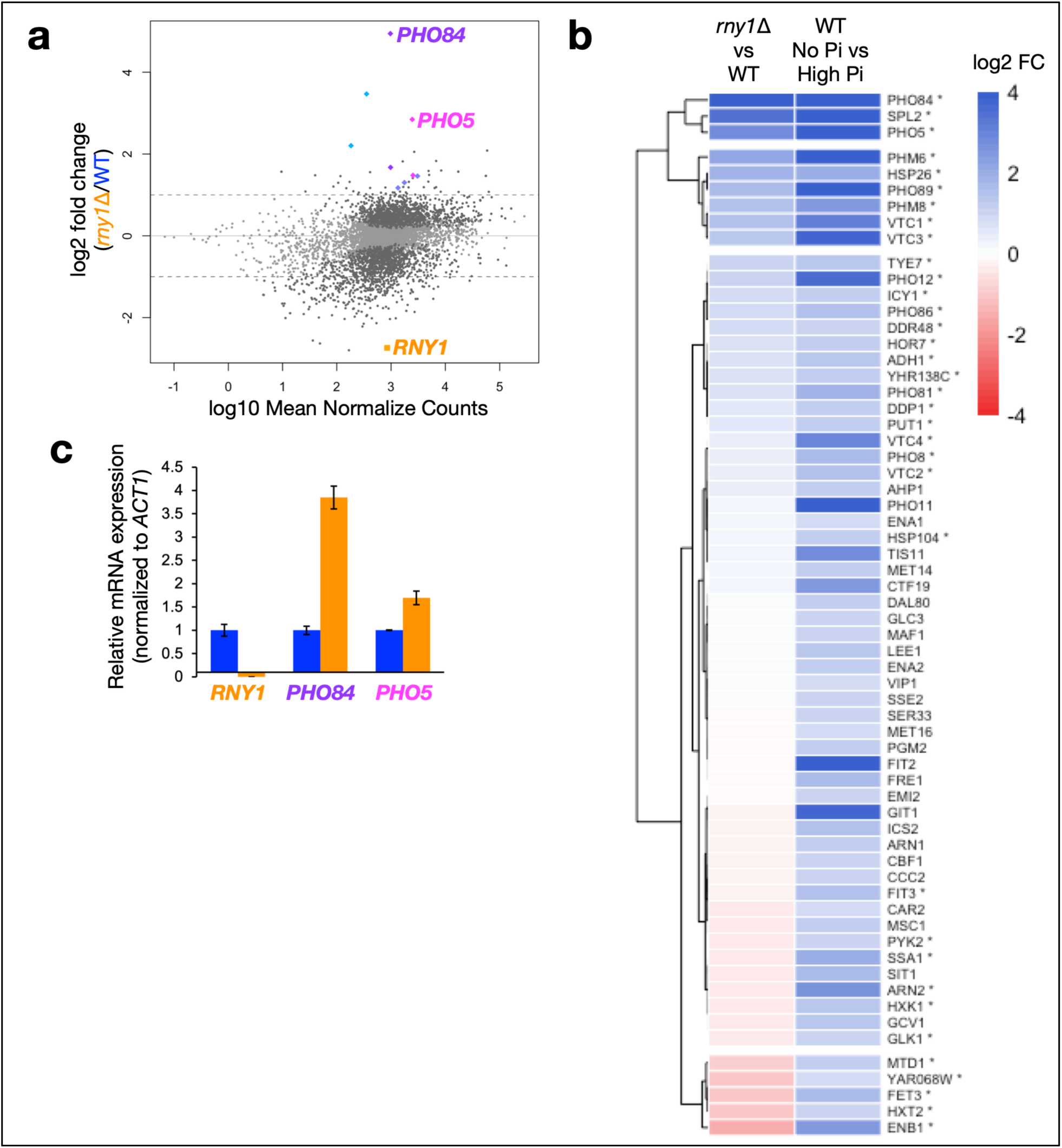
*Rny1* deficient cells exhibit gene expression patterns characteristic of phosphate starvation during growth in phosphate-rich media (A) Plot of log10 Mean Normalized Counts versus log2 fold change (*rny1Δ/*WT) displaying differential gene expression in Rny1-deficient (*rny1Δ*) strains versus wild-type during exponential growth in SC-LEU media. Significant changes in transcript abundance (≥2-fold change, *p*adj < 0.05) are highlighted in dark gray. The differential expression result for *RNY1* mRNA is plotted in orange. Selected Pho4-dependent target genes that significantly increase ≥2-fold in *rny1Δ*, specifically *PHO84* (purple) and *PHO5* (pink), are marked using color points. Data represent six poly(A)-selected RNA-seq libraries from three biological replicates per strain carrying an empty pRS415 plasmid. (B) Hierarchical clustering heatmap comparing a subset of the differential expression data from A against a published dataset of Pho4-regulated genes expressed during phosphate starvation (WT No Pi vs. High Pi; Zhou and O’Shea, 2011). The scale bar denotes log2 fold change of transcript level. Asterisks denote transcripts with significant gene expression changes in A (*p*adj < 0.05). (C) Relative mRNA expression level of *RNY1*, *PHO84*, and *PHO5*, normalized to *ACT1* mRNA levels in WT (blue) and *rny1Δ* (orange) strains carrying an empty pCM189 plasmid. Cells were grown to exponential phase in SC-URA. cDNA was quantified by SYBR Green RT-qPCR following phenol–chloroform extraction of RNA and DNase-treatment. Error bars show SEM of three technical replicates.

To validate the RNA-seq results, we created a *rny1Δ::KanMX* deletion in the WT BY4741 strain background. Total RNA was isolated from wild-type and *rny1Δ* cells grown in SC media until mid-log growth. Using reverse transcription followed by quantitative PCR, we evaluated the expression of *PHO84*, *PHO5* and *RNY1* by quantifying the cDNA from each gene and normalizing these values to the levels of *ACT1* cDNA. We observed a similar trend to our differential gene expression analysis, with the two *PHO* genes expressed at a higher level in *rny1Δ* than in wild-type cells. This suggests that *PHO* gene expression occurs more readily in yeast cells that lack RNase T2 function and are growing in P_i_-rich media.

The upregulation of the *PHO* pathway observed in *rny1Δ* mutants may directly be mediated by the phosphate responsive transcription factor Pho4. Transcription of the *PHO* genes is tightly controlled by regulators that directly bind to their promoters to either repress or enhance transcription. For instance, the promoter of *PHO84* contains established binding sites for the Pho4 transcriptional activator (Harbison *et al*. 2004). Conversely, Cbf1 is a constitutively expressed transcription repressor that binds to the same consensus sequence as Pho4 to repress *PHO* gene expression under specific conditions (Zhou and O’Shea 2011). Therefore, the induction of *PHO* genes in *rny1*Δ strains could stem from Pho4-dependent activation or a loss of Cbf1-mediated repression. Because the RNA-seq data showed no significant changes to *CBF1* mRNA levels (Fig. S1), we reasoned that Pho4-dependent activation was more likely to contribute to the *rny1Δ* phenotype.

To test this model, we constructed a reporter strain where the *PHO84* promoter drives *mCherry* expression (Fig. 2A). Under phosphate-limiting conditions, WT and *rny1*Δ strains exhibited robust mCherry expression as visualized by fluorescence microscopy (Fig. 2B, blue and orange). This expression is dependent on *PHO4* as *pho4*Δ cells harboring the reporter failed to exhibit mCherry expression and showed a dramatic reduction in integrated mCherry fluorescence density (Fig. 2B, purple). To test if the elevated *PHO* expression in *rny1*Δ mutants requires Pho4, we introduced the *pho4*Δ into *rny1*Δ strains carrying the *pPHO84-mCherry* reporter. The double mutant displayed an integrated fluorescence density distribution similar to the *pho4*Δ single mutant (Fig. 2B, brown); this result is further validated by quantitative RT-PCR showing a reduction *PHO84* transcripts in both *pho4Δ* and *pho4Δrny1Δ* strains when assessed under growth conditions described in Fig. 1 (Fig. S2). Taken together, this data demonstrates that aberrant *PHO* pathway activation in RNase T2-deficient strains is dependent on Pho4 activity rather than a loss of Cbf1-mediated repression.

**Figure 2:**
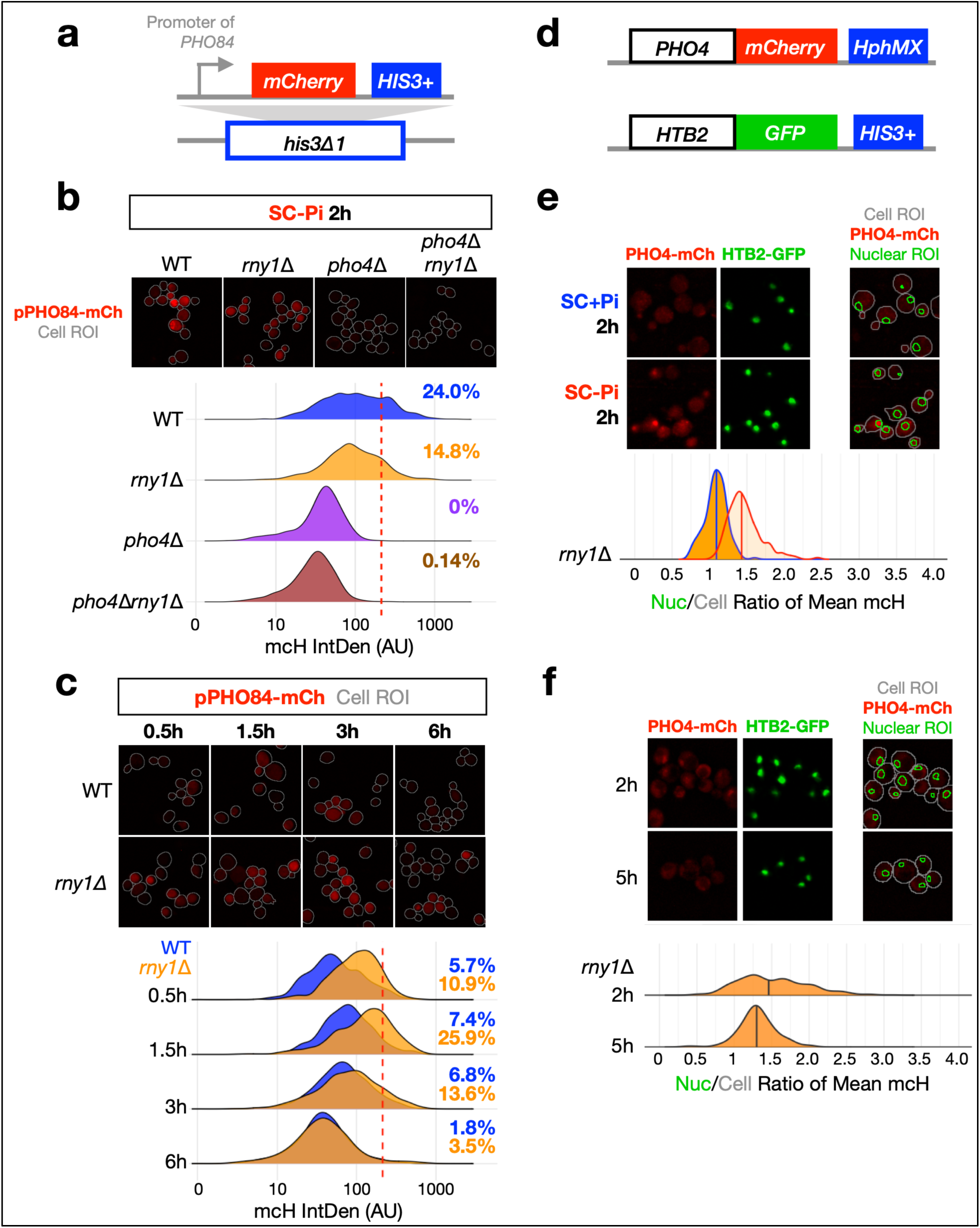
Aberrant Pho4 dependent transcriptional activation in RNase T2-deficient cells is observed upon dilution into phosphate rich media (A) Schematic of *pPHO84*-mCherry reporter gene used in panels B and C. (B) Top: Representative fluorescence microscopy images of cells back-diluted from overnight cultures into fresh SC media, grown to mid-exponential phase, and shifted into SC media without KH_2_PO_4_ (SC-P_i_) for 2 hrs. Bottom: Probability distribution function (PDF) plots of the integrated density of mCherry fluorescence for WT (blue), *rny1Δ* (orange), *pho4Δ* (purple), and *pho4Δ rny1Δ* (brown) cells. The red dashed line indicates the maximum reporter intensity observed in *pho4Δ* strains. Percentages indicate the fraction of cells exceeding this threshold. (C) Top: Representative fluorescence microscopy images of cells back-diluted from overnight cultures into fresh, SC media and sampled at 0.5h, 1.5hrs, 3hrs, and 6hrs post-dilution. Bottom: Corresponding PDFs of mCherry integrated density for WT (blue) and *rny1Δ* (orange) cells. Percentages indicate the fraction of cells exceeding the baseline threshold defined by the red dashed line in Panel B. (D) Schematic of the PHO4-mCherry fusion protein and the HTB2-GFP nuclear marker used for PHO4 localization assays in panels E and F. (E) Top: Representative images of *rny1Δ* strains expressing *PHO4-mCherry* and *HTB2-GFP* after 2 hrs of growth in either phosphate-rich medium (SC+P_i_, blue) or phosphate-depleted medium (SC-P_i_, red). The third column of panels represents mCherry fluorescence images with cell (gray) and nuclear (green) ROIs overlaid. Bottom: PDFs of the nuclear-to-cellular ratio of mean mCherry fluorescence intensity for *rny1Δ* cells grown in SC+P_i_ (blue line) or SC-P_i_ (red line). (F) Same as E except *rny1Δ* cells were back-diluted into fresh SC medium after overnight growth and imaged at 2hrs or 5hrs post-dilution.

We leveraged the *pPHO84-mCh*erry reporter to assess when *PHO* pathway activation occurred in *rny1Δ* mutants. WT and *rny1Δ* strains harboring the reporter were grown overnight in SC media, back-diluted to an OD_600_=0.2, and monitored for mCherry fluorescence over time during logarithmic growth (Fig. 2C). Shortly after back-dilution (0.5 h), *rny1*Δ cells exhibited noticeably higher mCherry expression than wild-type cells. This observation is consistent with our steady-state RNA-seq (Fig. 1A) and RT-qPCR (Fig. 1B) data and indicates that *rny1*Δ cells aberrantly activate the *PHO* pathway, despite being grown in a fresh and phosphate-rich media. This difference in *PHO* pathway activation was most evident 1.5 hrs post-dilution (Fig. 2C). At this time point, 25.9% of the *rny1Δ* population grown in rich media displayed mCherry fluorescence levels exceeding the maximum fluorescence seen in the *pho4Δ* strain grown in SC-P_i_ (red dashed line, Fig. 2B-C). By comparison, only 7.4% of wild-type cells shifted above this threshold. As growth continued to 6 hrs, the difference between *rny1*Δ and WT mCherry expression became less apparent (Fig. 2C, 6h). Collectively, these results suggest that aberrant *PHO* pathway activation in *rny1*Δ is more prominent at the initiation of growth in nutrient-rich media containing abundant extracellular phosphate.

To validate whether the aberrant Pho4-dependent transcription activation occurred more readily in *rny1*Δ cells shortly after back dilution, we monitored Pho4 localization in *rny1*Δ cells. To visualize the Pho4 localization, we expressed a *Pho4-mCherry* fusion alongside a nuclear marker, Histone H2B tagged with GFP (*HTB2-GFP*) (Fig. 2D). In phosphate-rich conditions, Pho4 is typically restricted to the cytoplasm, whereas under conditions of phosphate limitation, Pho4 translocates into the nucleus to drive transcription of the *PHO* genes. To ensure that the Pho4-mCherry fusion behaved as expected, the *rny1*Δ strain carrying both the *Pho4-mCherry* fusion and nuclear marker was grown overnight, back-diluted, and grown to OD_600_=0.5. Cells were then collected, washed and resuspended into either phosphate-rich SC media or phosphate-depleted SC-P_i_ media. After 2 hrs post depletion of phosphate, the *rny1*Δ cells exhibited clear Pho4 nuclear localization, as we observed increased mean mCherry fluorescence within HTB2-GFP defined regions of interests (ROIs) relative to the total cell ROI (Fig. 2E). This confirmed that the Pho4-mCherry fusion reflects the localization of Pho4 as expected.

We sought to determine if the aberrant expression of the *pPHO84-mCherry* reporter in Fig. 2C correlated with changes in the localization of Pho4. If *rny1*Δ cells aberrantly activate the *PHO* pathway during their initial growth in nutrient rich media, then we would expect to observe increased nuclear localization of Pho4 at early timepoints. When cultures were grown overnight in SC media, back diluted into fresh SC media, and observed 2 hrs post-dilution, a distinct population of cells exhibited higher nuclear localization of Pho4 (Fig. 2F) that is similar to that observed in cells grown in SC-P_i_ in Fig. 2E. By 5 hrs post-dilution, the cells displayed less nuclear mCherry localization, resembling the localization profile in SC+ P_i_ in Fig. 2E. This suggests that *rny1*Δ cells activated the *PHO* pathway more readily at the onset of growth in nutrient rich media.

Furthermore, these data suggest that RNase T2 function may be coupled with P_i_ metabolism. The activation of the *PHO* pathway in cells lacking RNase T2 function may be driven by an inability to maintain or rapidly rebuild internal phosphate pools. This could occur either because intracellular P_i_ stores are exhausted during overnight growth, necessitating a prolonged phase of rapid phosphate replenishment upon transfer to fresh media, or because these stores are utilized more rapidly than in wild-type cells to initiate growth.

### Loss of *Rny1* depletes intracellular polyphosphate stores and elevates total cellular RNA

To test the hypothesis that *rny1*Δ cells activate the *PHO* pathway because they are starved for phosphate, we examined internal phosphate pools. We expected intracellular phosphate levels to show depletion in the mutant as the *PHO* pathway is canonically activated when cells become starved for phosphate. WT and *rny1Δ* cells were grown overnight in SC media, back-diluted, and grown for an additional 1.5 hrs before being harvested, washed with water, and lysed. Total protein in the lysate was measured via a BCA assay and used for normalization. When free phosphate levels in the cell lysate were measured using a malachite green plate assay, a significant reduction in intracellular free phosphate levels was observed in *rny1*Δ strains (Fig. 3A). To measure total phosphate levels, we liberated phosphate bound within cellular macromolecules and polymers by boiling the lysate in sulfuric acid then quantified the phosphate released. Surprisingly, elevated total phosphate levels per µg of protein was observed in *rny1*Δ lysates when compared to WT lysates (Fig. 3A). These data suggest that while *rny1*Δ cells harbored more total phosphate than WT cells, they simultaneously harbored lower levels of available free phosphate and aberrantly activated the *PHO* pathway. Because RNA is an abundant, phosphate-rich macromolecule that comprises approximately 6–12% of cellular dry weight (Delgado *et al*. 2013), it represents a significant potential reservoir of intracellular P_i_. If Rny1-dependent RNA degradation were required to mobilize P_i_ within RNA, then *rny1*Δ strains would experience a deficit of intracellular phosphate because they could no longer utilize this P_i_ trapped in RNA for downstream anabolic reactions.

**Figure 3:**
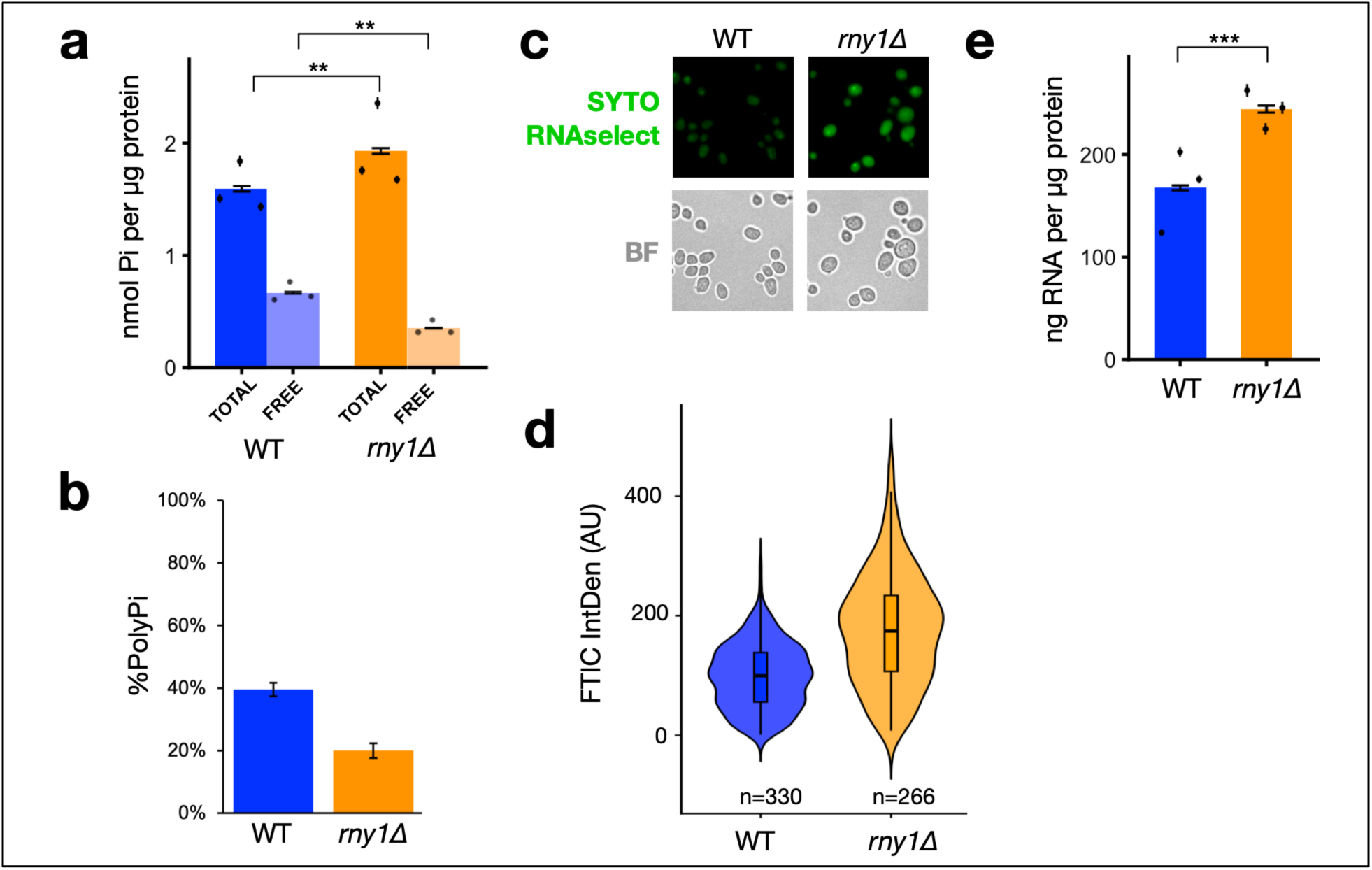
Intracellular phosphate is trapped in RNA in RNase T2 deficient cells (A) Total phosphate (TOTAL) and Free P_i_ (FREE) levels were measured in cell lysates from WT (blue) and *rny1Δ* (orange) cells after 1.5 hrs of growth in SC media after back-dilution. P_i_ amounts were quantified via malachite green assay and normalized to total protein content as determined by a BCA assay. The bar plot represents the average of 3 biological replicates, with individual biological data points plotted as black dots. Error bars for biological replicates represent the standard deviation (SD) of 3 technical replicates and error bars for the average represent the SD for 3 biological replicates. Double asterisks (**) indicate highly significant changes (0.001 < *p*-values < 0.01) as determined by a Welch’s t-test. (B) Polyphosphate (PolyP_i_) levels expressed as a percentage of the total isolated phosphate pool for WT (blue) and *rny1Δ* (orange) strains. The PolyP_i_ fraction was determined by calculating the ratio of free P_i_ released via exopolyphosphatase (PPX) digestion relative to the total phosphate content for an equivalent sample. Total phosphate content was determined by acid hydrolysis to release P_i_ from all phosphate-containing molecules. In both pools, free P_i_ was quantified using a malachite green phosphate assay. PolyPi was isolated from cells back diluted into fresh SC after overnight growth and grown to mid-log. Error bars represent the SD of three technical replicates. (C) Representative fluorescence and brightfield microscopy images of WT and *rny1Δ* strains grown overnight in nutrient-rich media and back-diluted into fresh SC media and grown to mid log. Cells were then stained with SYTO RNASelect, fixed with methanol and imaged. (D) Quantification of SYTO RNA fluorescence integrated density within Cellpose-generated regions of interest (ROIs) for wild-type (blue) and *rny1Δ* (orange) cells. Integrated density was calculated as the sum of pixel intensities within each ROI from background-corrected images. Violin and box plots show the distribution of single-cell measurements; *n* indicates the number of cells analyzed per strain. (E) Total RNA levels in cell lysates from WT (blue) and *rny1*Δ (orange) cells after 1.5 hrs of growth in SC media after back-dilution. RNA amounts were quantified using a fluorescence-based RNA quantification kit and normalized to total protein content as determined by BCA assay. Data is presented as described in A, except triple asterisks (***) indicate highly significant differences (0.0001 < *p*-value < 0.001) as determined by a Welch’s t-test.

This model yields two specific predictions: first, *rny1Δ* strains would utilize alternative intracellular phosphate stores to meet their anabolic demands, and second, *rny1Δ* mutants would accumulate undegraded RNA, which would account for the elevated total phosphate content observed in Fig. 3A.

To test if other intracellular stores of phosphate are utilized in *rny1*Δ cells, we examined the levels of polyphosphate (PolyP_i_), a key phosphate reserve in fungal cells. We quantified PolyP_i_ content in wild type and *rny1*Δ strains by isolating PolyP_i_ from mid-log cultures and performing an enzymatic digest with purified PPX, a specific exopolyphosphatase that releases P_i_ from PolyP_i_. Because the PolyP_i_ extraction contained significant amounts of co-purifying RNA, we normalized the assay by dividing the amount of free P_i_ released via PPX digestion by the total phosphate content of an equivalent sample subjected to complete heat and acid hydrolysis. For both measurements, free phosphate levels were quantified using malachite green assays. As shown in Fig. 3B, *rny1*Δ strains (orange) had significantly reduced PolyP_i_ levels relative to wild type (blue) cells. This result suggests that loss of *RNY1* function leads to a measurable reduction of stored phosphate reserves despite abundant extracellular phosphate. The reduction in polyphosphate levels provides a mechanistic basis for the activation of the *PHO* pathway in *rny1*Δ cells, as the *rny1*Δ cell may be sensing the scarcity of useable internal phosphate and consequently upregulating phosphate acquisition genes such as *PHO84* and *PHO5* to acquire more phosphate from the extracellular environment. Collectively, these findings raised the possibility that RNase T2 function contributes to maintaining intracellular P_i_ levels through RNA degradation and recycling of phosphate from RNase T2 decay products.

To test if the higher phosphate content observed in *rny1*Δ strains is due to increased total RNA in the cell, we measured total cellular RNA levels using fluorescent RNA-binding dyes. First, we utilized an *in vivo* approach by staining mid-log phase cells with the SYTO RNAselect dye. After 30 min of staining, cells were fixed with methanol and imaged (Fig. 3C). Mutant *rny1*Δ strains displayed significantly higher levels of SYTO RNAselect fluorescence than wild-type cells (Fig. 3C-D), even when accounting for the presence of strain-specific autofluorescence in *rny1*Δ cells (Fig. S3). To validate this *in vivo* finding, we generated cell lysates from *rny1*Δ and wild-type strains as in Fig. 3A and measured total RNA in the lysate using an Accublue RNA quantification kit, normalizing these values to total protein content. Consistent with data in Fig. 3C-D, we observed increased total RNA levels in *rny1*Δ when compared to WT (Fig. 3E). These findings, alongside previously published results (Huang *et al*. 2015, Minami *et al*. 2025), demonstrate that *rny1Δ* strains accumulate RNA, linking the activity of RNase T2 enzymes to the maintenance of intracellular phosphate homeostasis. If the inability to release phosphate stored in RNA directly drives this starvation signal, then restoring RNase T2-dependent RNA decay should reverse the phenotype.

### Genetic complementation with yeast or human RNase T2 suppresses aberrant PHO pathway activation

To test if the aberrant *PHO* pathway activation in *rny1*Δ strains can be reversed, we genetically reintroduced RNase T2 function into *rny1*Δ strains. To assess *PHO* pathway activation, we measured *PHO84* mRNA levels by RT-qPCR. As expected, *rny1*Δ cells exhibited significantly elevated *PHO84* mRNA levels relative to wild-type cells (Fig. 4A-B). When we expressed *Rny1* from a plasmid using its native promoter in *rny1*Δ cells, we observed significantly less *PHO84* expression, restoring mRNA levels similar to that observed in WT cells (Fig. 4A). This suggests that restoration of Rny1 activity rescues the aberrant *PHO* pathway activation. To test if the ability to degrade RNA was necessary for this rescue, we expressed a catalytically inactive form of the Rny1 (*rny1-ci*), where the catalytic histidine residues of the RNase T2 domain were mutated to alanine. When we measured *PHO84* mRNA levels in the *rny1*Δ + *rny1-ci* strains, we observed high levels of *PHO84* transcript that were comparable to what is observed in *rny1*Δ strains without a rescue (Fig. 4A). This indicates that the RNA decay activity of Rny1 is strictly required for preventing activation of *PHO84*. While the *S. cerevisiae Rny1* contains a unique C-terminal domain, the core RNase T2 domain is highly conserved from yeast to humans. To test if this conserved core activity is sufficient for phosphate homeostasis, we created a plasmid that expresses the human RNase T2 enzyme fused to a yeast vacuolar signal sequence and transformed it into *rny1*Δ cells. We observed that the human RNase T2 homolog successfully rescued *PHO84* expression to levels observed in wild type cells (Fig. 4B). This complementation further supports the idea that degraded RNA can act as a crucial source of P_i_ required for phosphate homeostasis, and suggests a potentially conserved role for RNase T2 enzymes in phosphate recycling.

**Figure 4:**
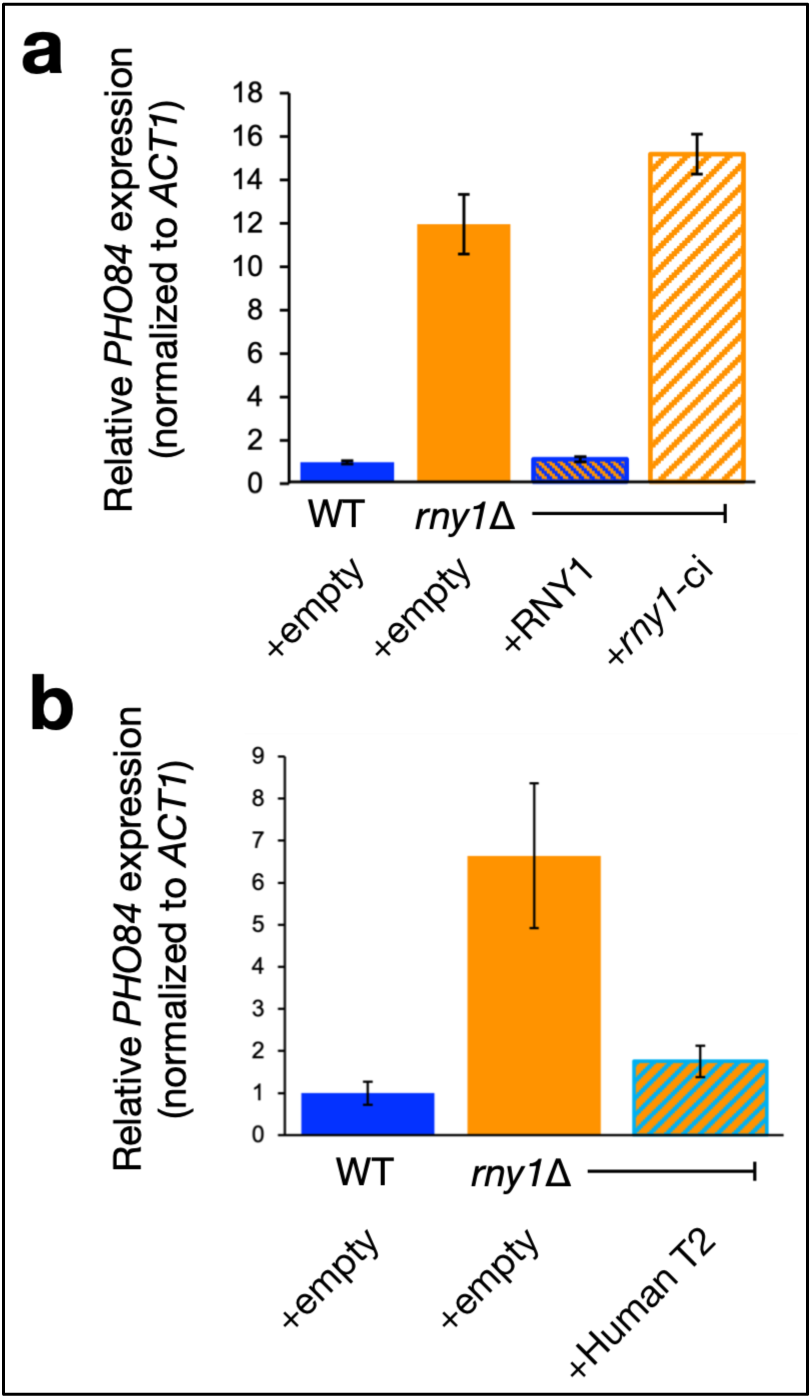
Aberrant PHO pathway activation in *rny1*Δ is rescued by the RNA decay activity of RNase T2 (A) Relative *PHO84* mRNA expression normalized to *ACT1* in WT (blue), *rny1*Δ (orange) and *rny1*Δ mutant strains complemented with plasmids expressing yeast *RNase T2* constructs (orange outlined). Total RNA was isolated from cultures backdiluted from overnight growth in selective SC-LEU media and harvested at OD600≈0.6; mRNA levels quantified as in Fig. 1B. Error bars represent SEM of three technical replicates. *rny1Δ* strains were complemented with wild-type *RNY1* or a catalytically inactive mutant (*rny1*) carrying H→A substitutions in catalytic residues required for RNA decay. (B) Same as in A, except strains were grown in selective SC–URA media. The *rny1Δ* strain was complemented with human RNase T2 fused to a yeast signal sequence.

### *RNY1* mRNA is upregulated in response to phosphate limitation

We propose that RNA degradation by RNase T2 provides a critical source of intracellular phosphate for degraded RNA. We hypothesize that this activity becomes vital during extracellular phosphate limitation, when cells have a limited amount of free P_i_ to synthesize essential macromolecules like phospholipids, DNA, and ATP, and must recycle or utilize internal P_i_ stores. Consistent with this model, phosphate limitation upregulates plant RNase T2 enzymes (Bariola *et al*. 1994, Gho *et al*. 2020). While *RNY1* expression and vacuolar localization have primarily been studied under nitrogen depletion (Huang *et al*. 2015, Minami *et al*. 2025), the promoter region of *RNY1* notably contains putative Pho4 consensus binding sites (Zhou and O’Shea 2011). To test the hypothesis in yeast, we examined *RNY1* and *PHO84* expression by RT-qPCR under several different nutrient-deprived growth conditions. To evaluate *RNY1* expression, a wild-type culture was grown to mid-log phase and shifted for two hrs into SC media (SC), media lacking nitrogen (SD-N), or media lacking phosphate (SC-P_i_) (Fig. 5A). As expected, *RNY1* expression was low in cells grown in SC media but induced in cells grown in SD-N (Fig. 5C). Conversely, we did not observe strong *PHO84* expression in SC or SD-N (Fig. 5B). However, in cells grown in SC-P_i_, we observed strong *PHO84* expression.

**Figure 5:**
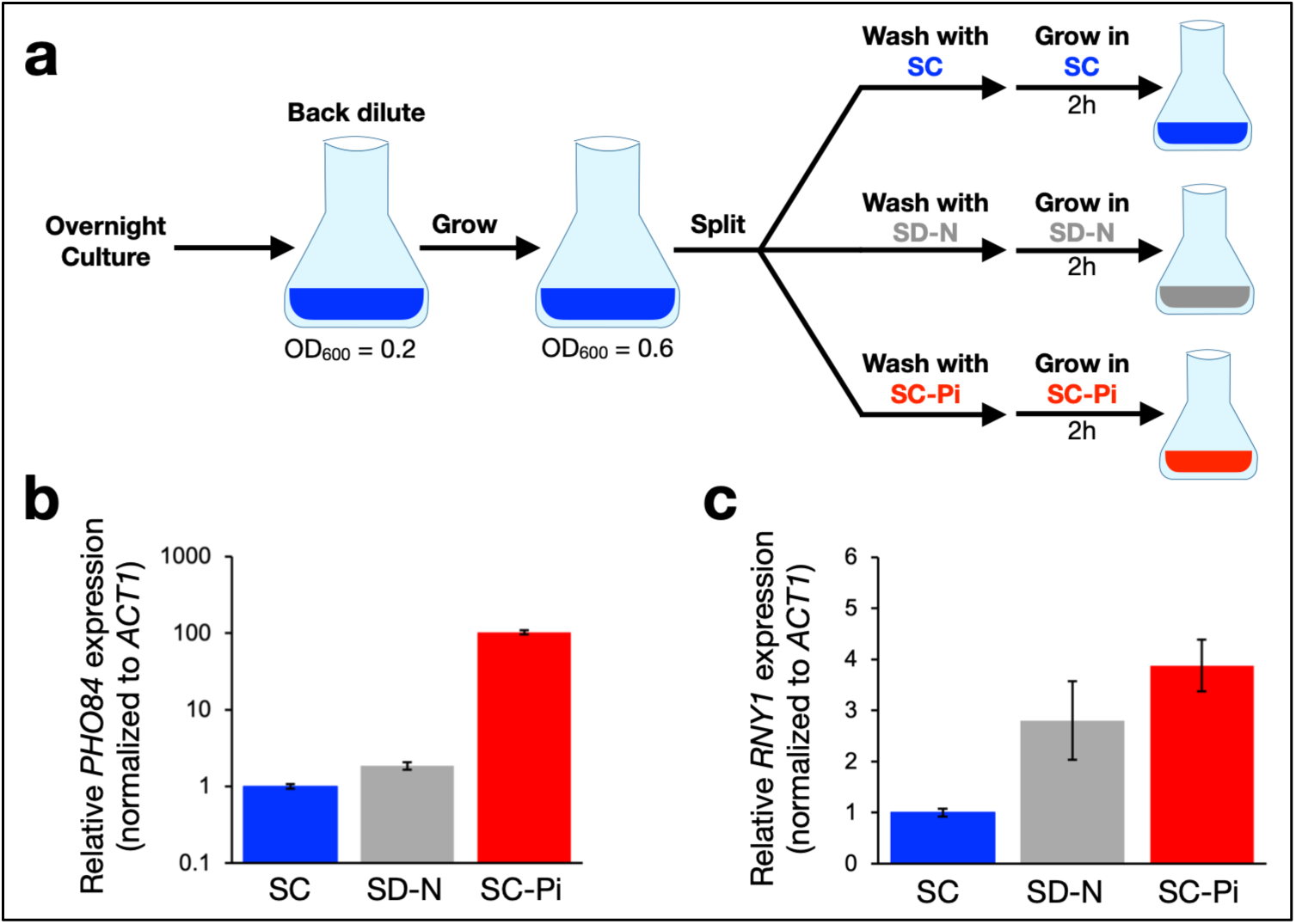
Rny1 is expressed under phosphate limiting conditions (A) Schematic of experimental design. Cells were grown overnight in SC, back-diluted, and grown to mid-log phase (OD600 ≈ 0.6), then split, washed, and shifted into SC, nitrogen-depleted SC (SD–N), or phosphate-depleted SC (SC–Pi) for 2 hrs. (B-C) Relative *PHO84 (B)* or *RNY1 (C)* mRNA expression normalized to *ACT1* in WT strains shifted from SC into SC media (blue), nitrogen-depleted SC media (gray), or phosphate-depleted SC media (red). RNA extraction and RT-qPCR as in Fig. 1B. Error bars represen SEM of three technical replicates.

Strikingly, *RNY1* expression was also upregulated in SC-P_i_, reaching similar levels observed in SD-N (Fig. 5C). These results demonstrate that *RNY1* is expressed in response to phosphate limitation, further supporting our model that RNase T2-mediated RNA degradation can serve as a metabolic pathway to liberate phosphate under phosphate-limiting growth conditions.

## Discussion

Our work demonstrates that the *S. cerevisiae* RNase T2 enzyme, Rny1, is necessary for phosphate homeostasis, where it functions to free inorganic phosphate (P_i_) from degraded RNA (Fig. 6). Cells lacking RNase T2 activity exhibit aberrant Pho4-dependent activation of the PHO pathway even under high phosphate growth conditions. This phenotype, combined with our observations that *rny1Δ* cells accumulate total RNA and exhibit higher total phosphate content, indicates that phosphate can become metabolically trapped in RNA normally degraded by RNase T2. In line with this model, we also observe reduced polyphosphate stores in *rny1*Δ cells. Together, these findings position RNA turnover by RNase T2 enzymes as a central mechanism in maintaining phosphate homeostasis. Rather than relying solely on extracellular phosphate transport or vacuolar mobilization of PolyP_i_, cells can dynamically tap the intracellular RNA pool. This work positions RNA as a reservoir of inorganic phosphate that promotes cellular homeostasis.

**Figure 6:**
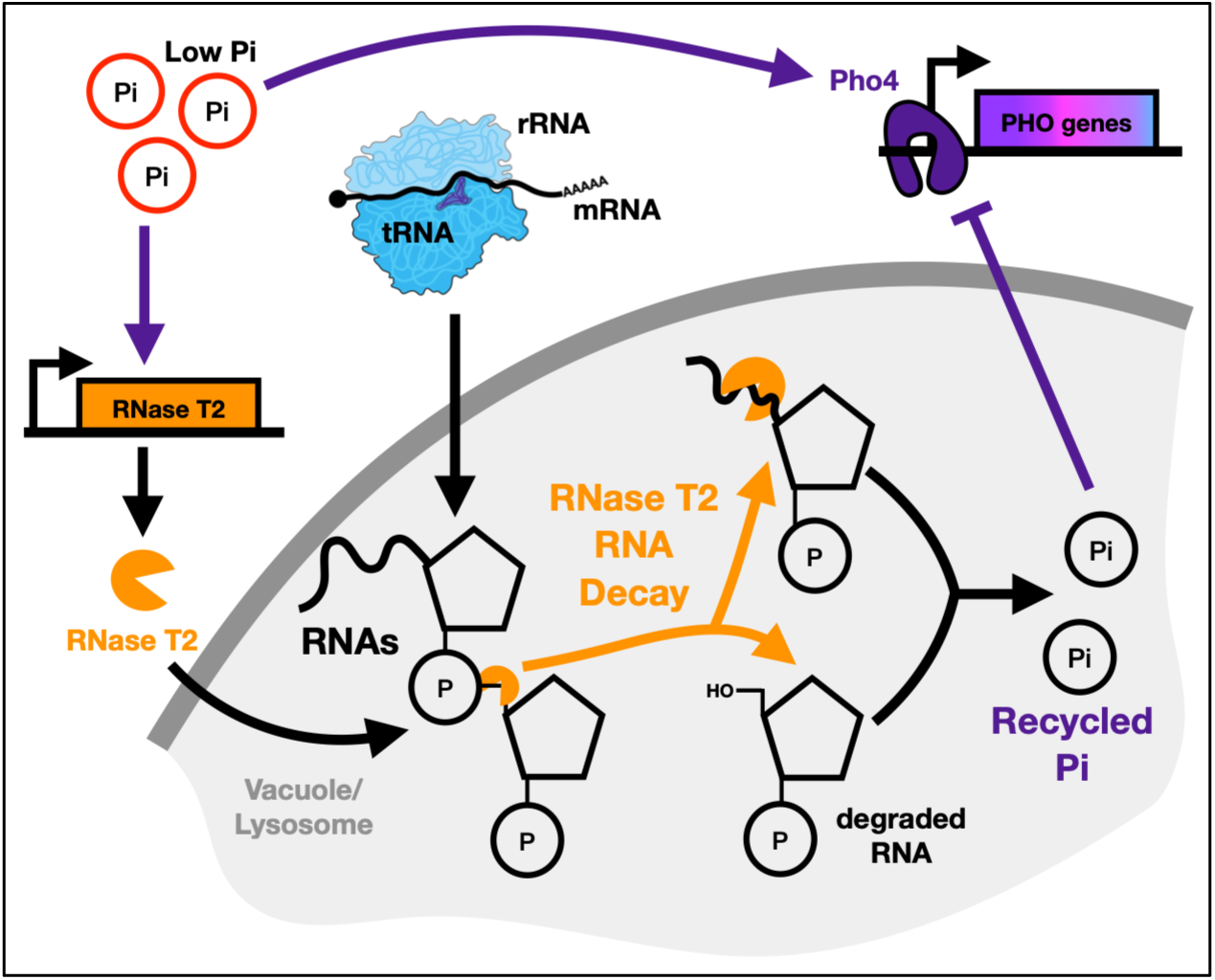
Proposed model of RNase T2-mediated Phosphate Homeostasis Under low inorganic phosphate (Pi) growth conditions, the transcription factor Pho4 is activated to induce the expression of *PHO* genes. Concurrently, RNase T2 expression is upregulated along with other factors necessary for autophagic RNA degradation within the vacuole/lysosome. RNase T2 degrades these imported RNAs, thus recycling Pi into the cytoplasm. This RNAderived Pi subsequently inhibits Pho4 activation of *PHO* genes, establishing a negative feedback loop to regulate phosphate homeostasis.

This dynamic recycling of RNA for an essential metabolite highlights an elegant metabolic strategy, particularly in that it exploits RNA’s natural abundance and rapid turnover, which are inherent to most cells. RNA primarily serves to transfer genetic information and drive protein synthesis, with ribosomal RNA (rRNA) constituting most of the cellular pool. This large cellular investment into generating RNA also represents the production of an abundant and resource-rich intracellular store of phosphate. Consequently, viewing RNA as a phosphate reservoir reveals a straightforward negative feedback loop: under cellular stress, cells harvest inorganic phosphate from RNA while simultaneously downregulating translation by degrading rRNA, tRNA, and mRNA pools. Supporting this model, rRNA is a conserved target of RNase T2 enzymes across diverse organisms, ranging from plants (Hillwig *et al*. 2011) to fish (Haud *et al*. 2011). Because translation is one of the most energetically demanding processes in the cell (Warner 1999, Lynch and Marinov 2015), coupling targeted RNA decay via Rny1 to phosphate recycling allows the cell to conserve energy during P_i_ limitation. Therefore, RNA decay by RNase T2 enzymes may be a finely tuned metabolic adaptation to survive the limitation of one of the most critical molecules of life: inorganic phosphate.

This mechanism reflects a broad evolutionary conservation of phosphate recycling strategies across eukaryotes. In plants, including *Arabidopsis thaliana* (Bariola *et al*. 1994, Hillwig *et al*. 2011), tomato (Löffler *et al*. 1992), and rice (Gho *et al*. 2020), a general link between RNase T2 upregulation and phosphate limitation has been observed. However, while these plant systems highlight a conserved survival response, our work in yeast defines the precise physiological and regulatory consequences of this pathway. By demonstrating that the loss of *Rny1* leads to the over-accumulation of total RNA, depleted polyphosphate stores, and an aberrant Pho4-dependent activation of the *PHO* pathway, we establish that RNase T2-mediated RNA turnover is not merely a passive stress response, but a finely tuned homeostatic feedback loop. Consistent with this model, our data mining of an RNA-seq dataset from the distantly related budding yeast *Candida albicans* revealed that the RNase T2 family member *RBT7* is also strongly upregulated during phosphate starvation (Ikeh *et al*. 2016).

While the conservation of this system across species is clear, our finding that *RNY1* expression is upregulated in SC-P_i_ media is unexpected, given that transcriptional responses to phosphate limitation have been extensively profiled in budding yeast (Mouillon and Persson 2006, Zhou and O’Shea 2011). Across multiple published transcriptomic datasets that we examined, *RNY1* was never identified as upregulated during phosphate limitation, nor has it been traditionally classified as a *PHO* gene. Interestingly, despite the *RNY1* promoter containing a canonical Pho4 consensus motif, Pho4 ChIP-seq data do not show enriched Pho4 binding under low-or no-phosphate growth conditions relative to high-phosphate controls. Instead, our data mining revealed an increase in Cbf1 binding at the *RNY1* promoter specifically under no-phosphate conditions (Zhou and O’Shea 2011). Therefore, characterizing the transcriptional networks and activation dynamics governing *RNY1* expression during phosphate limitation, and defining how they intersect with other nutrient stress responses such as nitrogen starvation, presents an important avenue for future research.

Lastly, these findings raise a critical mechanistic question: how does the loss of Rny1-mediated RNA decay trigger the aberrant activation of the *PHO* pathway? A key clue lies in the metabolic fate of degraded RNA during growth under nitrogen starvation. Work by Huang et al. (2015) demonstrated that during bulk RNA turnover by autophagy, the resulting 3’-nucleotide monophosphates are further processed to release the 3’ phosphate group. The remaining nucleoside and base byproducts are then actively broken down and excreted from the cell. This observation strongly implies that the primary metabolic objective of Rny1 activity is the selective acquisition of inorganic phosphate. When *Rny1* is deleted, this essential Pi pool remains trapped within RNA in the vacuole, creating an internal phosphate deficit that likely explains the *rny1Δ* phenotypes observed in our study. The drop in PolyP_i_ likely occurs because the cell must continuously hydrolyze existing stores to meet cytoplasmic demand, or because a localized P_i_ supply chain between vacuolar Rny1 and the PolyP_i_-synthesizing VTC complex is severed. This internal P_i_ crisis provides a direct mechanism for *PHO* pathway activation. A compelling genetic precedent exists in purine metabolism, where deletion of the adenine deaminase *AAH1* derepresses the *PHO* pathway under high-phosphate conditions due to an internal metabolic imbalance (Choi *et al*. 2017). We hypothesize that localized or systemic P_i_ depletion in the *rny1Δ* mutant causes a similar intracellular disruption. Ultimately, by failing to free phosphate from RNA, the cell generates an internal starvation signal, activating the regulatory network to constitutively activate Pho4 even when extracellular P_i_ is abundant.

## Data Availability Statement

Strains and plasmids generated in this study are available upon request. The raw and processed RNA-sequencing datasets generated and analyzed during this study have been deposited in the NCBI Gene Expression Omnibus (GEO) repository and are accessible under accession number GSE338688.

## Acknowledgements

The authors thank the students and staff associated with the Spring 2022 MB 365: Genomics lab course at Colorado College for generating the poly(A)-selected libraries used to produce the data in Figure 1. We thank the Technical Director, Carrie Moon, for logistical support, Sara Hanson for course materials, and students A. Fan, S. Yam, A. Mat, and M. Ort for library preparation. We also thank previous undergraduates in the Garcia Lab for generating strains and testing protocols used in this study: Hayden Low, Rachael Martino, Jocelyn Zuckerman, Rufino Driscoll, and Connor Pepin.

We thank Hiten Madhani and Eva Balog for critically reviewing the manuscript and for helpful discussions.

Sequencing was performed at the University of California, San Francisco (UCSF) Center for Advanced Technology (CAT). This facility is supported by UCSF Program in Breakthrough Biomedical Research (PBBR), the Resource Allocation Program (RRP) Institutional Matching Instrument Award (IMIA), and the National Institutes of Health (NIH) under grant number 1S10OD028511-01.

We gratefully acknowledge the University of New England Histology and Imaging Core for providing access to microscopy instrumentation and technical support.

